# Easydecon: Efficient Cell Type Mapping for High-Definition Spatial Transcriptomic Data

**DOI:** 10.1101/2025.04.08.647764

**Authors:** Sinan U. Umu, Victoria T. Karlsen, Espen S. Bækkevold, Frode Lars Jahnsen, Diana Domanska

## Abstract

The emergence of high-resolution spatial transcriptomics platforms, such as VisiumHD, has enabled transcriptome-wide spatial profiling at near-single-cell resolution. However, existing analysis tools often lack scalability or compatibility with this new resolution, limiting their utility for multimodal cell type analysis. We present Easydecon, a lightweight and modular computational framework for spatial transcriptomics analysis using marker genes from single-cell RNA sequencing datasets. Easydecon uses a two-phase strategy, which firstly detects expression hotspots and then refines cell type assignments with similarity-based methods. We demonstrate its efficacy by resolving cell type subsets with high accuracy. Easydecon supports integration with segmentation tools and outperforms established methods in speed, usability and cell type recovery.

## Introduction

Biological tissues are composed of various cell types and extracellular elements, and their functionality is determined by both these components and their spatial arrangement within the tissue [1]. Novel imaging and molecular profiling tools allow us to study these tissues in more detail than before, giving us a better understanding of their composition and organization. Spatial transcriptomics (ST) uses RNA sequencing (RNA-seq) to investigate cells and their gene expression patterns within their spatial context [2]. It has revolutionized research in areas such as developmental biology, cancer, and neuroscience by enabling integration of molecular data with spatial organization, overcoming limitations of scRNA-seq alone [3]. In recent years, we have seen major developments in ST technologies such as resolution, multiplexing capacity, compatibility with clinical samples, enabling more detailed, accurate, and scalable analysis, which allow us to better understand spatial patterns among various diseases and tissues [4–7].

The Visium platform from 10x Genomics has been one of the most popular among the ST platforms. This allows spatial resolution of target tissue resolved in 55 μm bins. The platform has already enabled better understanding of the heterogeneous nature of cancer and tumor microenvironment (TME) [8,9]. However, the analyses and single-cell data integration of ST usually require computationally expensive deconvolutional tools for seamless integration with other modalities and for fine-grained spatial resolution [10]. The new version of the 10x Visium platform, VisiumHD, achieves near single-cell resolution with smaller and gapless spots i.e. bins. This partly solves the problem of the old Visium technology, where the same spots contain multiple cells, and may ease the requirement of complex deconvolutional algorithms. The new platform does this by offering different bin sizes with the default 8 µm resolution having more than half a million squared bins on a single slide. It is also possible to use 2 µm (offering millions of squared 2×2 µm spots) or 16 µm sized bins, which is a tremendous improvement compared to the previous version [11].

However, the efficient analysis of VisiumHD data is not straightforward as most of the tools have been developed considering the previous version or other ST platforms. Many deconvolutional tools are also not working efficiently on very large VisiumHD objects. This poses challenges for researchers who want to do multimodal cell type analysis using data coming from various scRNA-seq platforms. Fortunately, some of the frameworks like SpatialData [12] operate effectively on different ST technologies including VisiumHD which creates a strong basis for analysis. There are also recent solutions for analysis like Bin2cell [13], which utilizes 2 µm bins (or squares) to identify segmented cells on the H&E image, or Enact workflow, which allows segmentation followed by multiple cell type annotation methods [14]. Yet, cell type analysis is still not streamlined to utilize any scRNA-seq data and requires additional complex bioinformatics solutions.

In this study, we describe Easydecon, a novel workflow to analyze high definition spatial datasets. Since visualization of distinct cell types and markers are still one of the major challenges, Easydecon utilizes lightweight algorithms and regression-based estimation for marker visualization, single-cell cluster transfer and deconvolution. Easydecon uses the latest bioinformatics packages, compatible with VisiumHD, and enables fast analyses of high-definition ST datasets. It is a Python package, available at PyPi, with minimal dependencies and any scRNA-seq dataset coming from different single-cell technologies (i.e. 10x Genomics, Parse biosciences etc.) can be used. We tested Easydecon on publicly available ST-datasets from colorectal cancer (CRC), normal colon, tonsil and standard annotations as a showcase. We demonstrated how it can be used effectively and easily.

Easydecon is available at GitHub (github.com/sinanugur/easydecon) and PyPi.

## Materials and Methods

### Easydecon Analytical Framework

Easydecon follows a two-step approach. The first step focuses on identifying spatially informative marker gene expression patterns based on differential expression analyses from scRNA-seq or from another ST dataset. These markers are then assessed within the VisiumHD dataset by aggregating expression levels in spatial bins using selected aggregation methods (e.g., sum, mean, median). To identify significant marker gene expression spots, Easydecon uses filtering algorithms like quantile-based, permutation-based or negative binomial approaches to separate meaningful expression signals from background noise. This helps refine marker-gene expression profiles for each spatial bin. Next, Easydecon assigns cell type labels to these spatial bins by calculating similarity scores between spatial bin expression profiles and predefined marker gene profiles from scRNA-seq data. It does this by calculating similarity scores using different metrics, such as correlation, cosine similarity or weighted Jaccard. This step ensures that spatial bins are efficiently matched to distinct cell types or clusters, effectively integrating scRNA-seq data into spatial analysis. The results can be merged back into the VisiumHD dataset, enabling streamlined visualization, downstream analysis and interpretation.

### Detection of Expression Hotspots

Easydecon employs a refined permutation test to robustly identify significant expression hotspots. This method generates an empirical null distribution by repeatedly aggregating random gene sets drawn from the most variable gene pool (e.g., top 30% of variable genes). The null distribution is generated from spatial bins exhibiting moderate expression levels, defined by excluding bins at the extreme quantiles (default 10% highest and lowest expression bins). To enhance statistical robustness, permutation subsampling is divided into multiple subsets, each with a specified number of permutations (default total 5000 permutations). Null distributions from these subsets are combined, ensuring comprehensive representation of potential background variation.

Two thresholding strategies are supported:

*Parametric Thresholding:* Fits either gamma or exponential distributions to the empirical null, deriving a threshold from the inverse cumulative distribution function at a predefined significance level (α).

*Empirical Thresholding:* Directly calculates the significance threshold as the (1 - α) quantile of the permutation-based null distribution.

Moreover, filtering can be performed using quantiles, providing an alternative approach for refining expression hotspots.

### Similarity Algorithms

#### Weighted Jaccard Similarity

This extends the classic Jaccard similarity by incorporating weights to reflect gene importance or expression levels. This method is particularly suitable for comparing expression profiles between spatial bins and reference marker gene sets.

The weighted jaccard similarity *J_w_* between two gene sets *A* (reference markers) and *B* (spatial bin expression) is defined as follows:

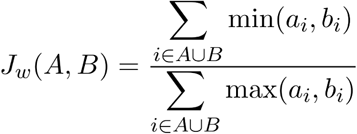

where:

- *A U B* denotes the union of genes from both sets.
- *a_i_*and *b_i_* represent the weights associated with the gene *i* in the reference marker set and the spatial bin, respectively.

Weights for genes can be assigned using two distinct strategies:

- Pre-calculated Weights: If available, predefined weights derived from differential expression analyses (e.g., log fold-changes) are normalized and directly utilized as gene weights.

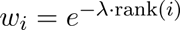

- ● Rank-based Exponential Decay: If predefined weights are unavailable, gene weights are assigned based on their rank positions within the marker gene list using an exponential decay function:

where *rank(i)* denotes the rank position of gene *i*, and the parameter *λ* controls the decay rate, emphasizing genes ranked higher in significance. The resulting Weighted Jaccard index provides a robust measure of similarity that captures both the presence of genes and their relative importance or expression strength, enhancing the accuracy of spatial cluster assignments.

#### Other Similarity Metrics

We also utilize other similarity metrics. Szymkiewicz–Simpson similarity also known as overlap similarity. It evaluates similarity based on the intersection relative to the smaller set and emphasizes complete or nearly complete overlap of gene sets. Cosine similarity based on the cosine of the angle between two gene-expression vectors, emphasizing directional concordance. Here we use the normalized spatial expression and fold changes or scores. Jaccard similarity measures the overlap between two gene sets relative to their union. Since we measure similarity in each bin spot separately, similarity metrics are robust to enable a direct similarity comparison among cell clusters.

### Cluster Assignment Strategies

Clusters are assigned to spatial bins based on calculated similarity metrics. Two primary approaches are supported:

Direct Maximum Similarity (“max”): Each bin is assigned to the cluster with the highest similarity or proportion score.

Hybrid Adaptive Assignment (“hybrid”): A softmax-based probabilistic approach is used, incorporating both similarity magnitude and relative distribution across clusters to robustly assign bins, especially handling ambiguous cases.

The cluster assignments are merged back into the dataset, enabling visualization and further spatial analyses.

### Differential Expression Analyses

Easydecon relies on DE tables. The marker genes for colon atlas were identified from the single-cell dataset using the Scanpy framework (v1.11.2). Standard preprocessing steps were applied, including library size normalization, log-transformation, and highly variable gene selection using default Scanpy parameters. The analysis was performed using the Wilcoxon rank-sum test, as implemented in Scanpy, comparing each annotated cell type or cluster against all others. Only genes with adjusted p-values below 0.05 and positive log fold changes (0.25) were retained as marker candidates. For each cell type, the top genes were included to construct the DE table used by Easydecon. This table serves as the primary source of marker gene signatures for downstream similarity and cell type analyses.

For the tonsil atlas analysis, we used the Bioconductor HCATonsilData single-cell reference dataset [15], which was downloaded and processed in R (v4.4.3). The differential expression analysis was performed using the scran package (v1.34.0) with default normalization and modeling parameters. Marker genes for each annotated cell type were identified based on the default pairwise differential expression workflow in findMarkers, which applies empirical Bayes moderation to detect genes with consistent upregulation across cells of the same type. The resulting marker gene list was compiled into the differential expression table used for Easydecon analysis of the tonsil VisiumHD dataset. We kept the top 20 genes based on the summary area under to curve (AUC) values.

### Parameters Used for Easydecon and Methods Benchmarking

We used the default Easydecon parameters for all tests: the top 60 marker genes per group based on either the Scanpy score or the scran AUC score, a phase-1 filtering threshold of p < 0.01, and wJaccard similarity computed from log fold changes. We also used log fold changes to estimate cell type proportions (). The VisiumHD samples were processed and log-normalized after initial QC (min_cells = 10, min_genes = 10), with mitochondrial genes removed. All benchmarked tools were run with their respective default settings. The single-cell datasets were already preprocessed and no additional modifications were applied other than normalization when required, using identical parameters.

## Results

### Easydecon workflow allows fast ST analysis

Easydecon is designed to operate on the VisiumHD platform optimally for 8 µm resolution bins while the platform also offers 2 µm and 16 µm resolutions. Easydecon utilizes permutation tests, similarity metrics and bin decomposition modelling to transfer cell type labels from scRNA-seq datasets to high resolution ST data or across different ST samples. The analysis workflow starts from tables representing differentially expressed (DE) markers, in CSV or Excel format. By relying on pre-computed DE results, Easydecon ensures efficient and reliable label transfer while maintaining high computational performance. Furthermore, it supports the use of selected marker genes for different cell types, which can serve as pre-filtering markers or can be directly mapped onto spatial locations for basic visualization.

This design allows researchers to easily integrate any publicly available or own scRNA-seq data with their high definition ST datasets, making it straightforward to identify distinct cell types, cell type proportions and spatial locations without relying on complex, computationally expensive deconvolution algorithms (Fig. 1).

**Figure 1.**
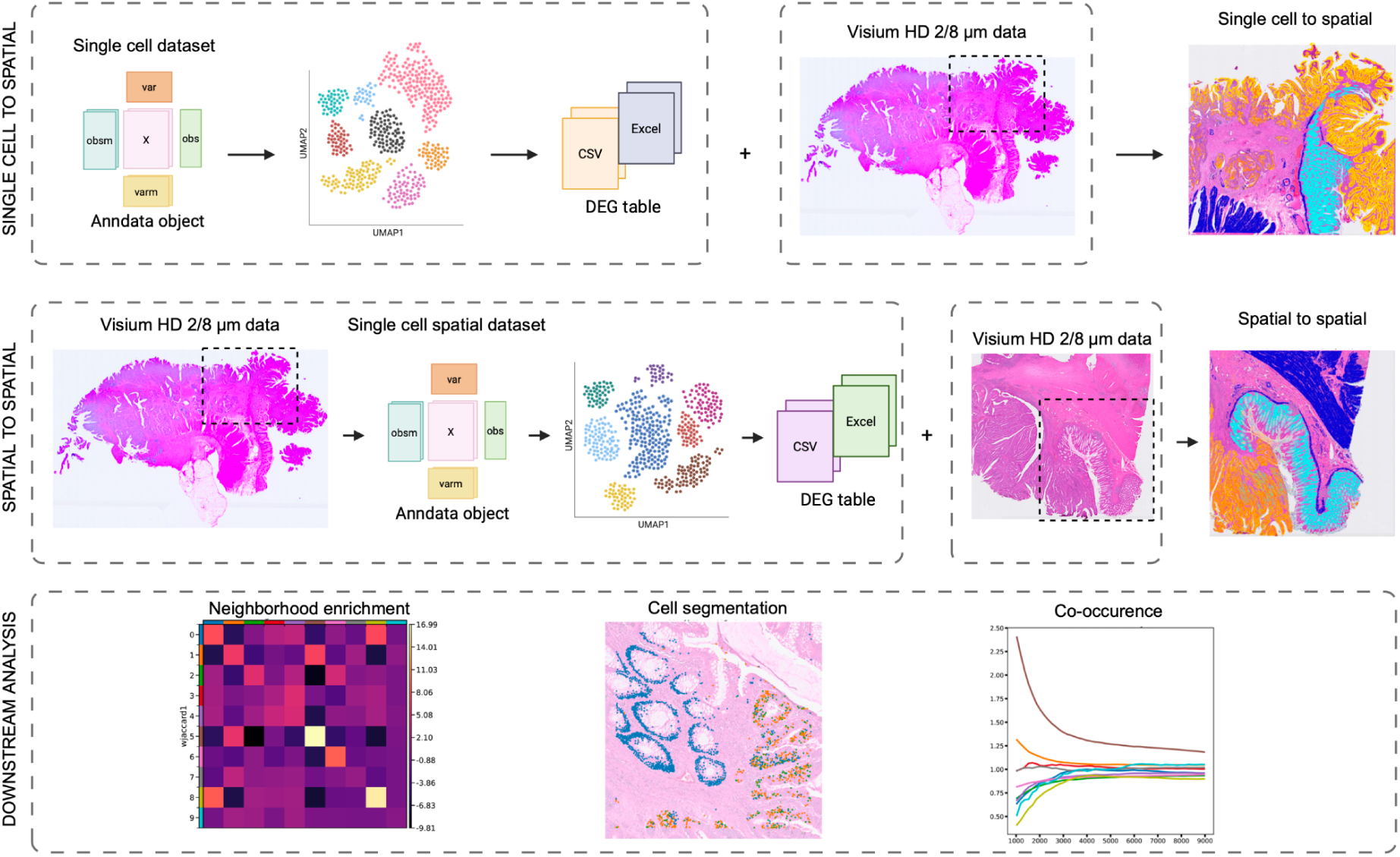
Overview of the Easydecon workflow shows the use cases. Easydecon supports both single-cell-to-spatial and spatial-to-spatial data mapping, as well as cell type proportion estimation, using either simple DE tables or single-cell objects in Anndata or Seurat formats. It also integrates seamlessly with cell segmentation tools such as Bin2cell. After annotation, users can perform various downstream analyses at either the bin or cell level, including neighborhood enrichment, quantification and co-occurrence assessment.

### Easydecon enables flexible cell type transfer from single-cell datasets using DE markers

As a general and robust test case for Easydecon, we acquired a publicly available single-cell CRC atlas dataset [16] and a VisiumHD CRC sample provided by 10X Genomics (Sample P2). Easydecon is designed to leverage scRNA-seq data without requiring the large single-cell object. Easydecon employs a two-step approach.

In the first step, global markers (e.g. major cell lineages markers or top-level markers) are used to detect expression hotspots, meaning the most likely positions of target cells among bins. These global markers can be manually selected based on prior knowledge and existing literature or directly extracted from the DE tables. In our test case, we used seven top-level clusters: epithelial (epi) cells, stromal (strom) cells, T-NK-ILC (TNKILC) cells, plasma cells, mast cells, B cells, and myeloid cells from the acquired CRC atlas dataset [16]. We retained the 60 highest-scoring markers for each cluster (Fig. 2a shows the top five). In this low-granularity step, bins with the strongest marker expression are flagged as the most likely locations of the corresponding top-level clusters (Fig. 2b), while Easydecon discards bins with weak or absent expressions.

**Figure 2.**
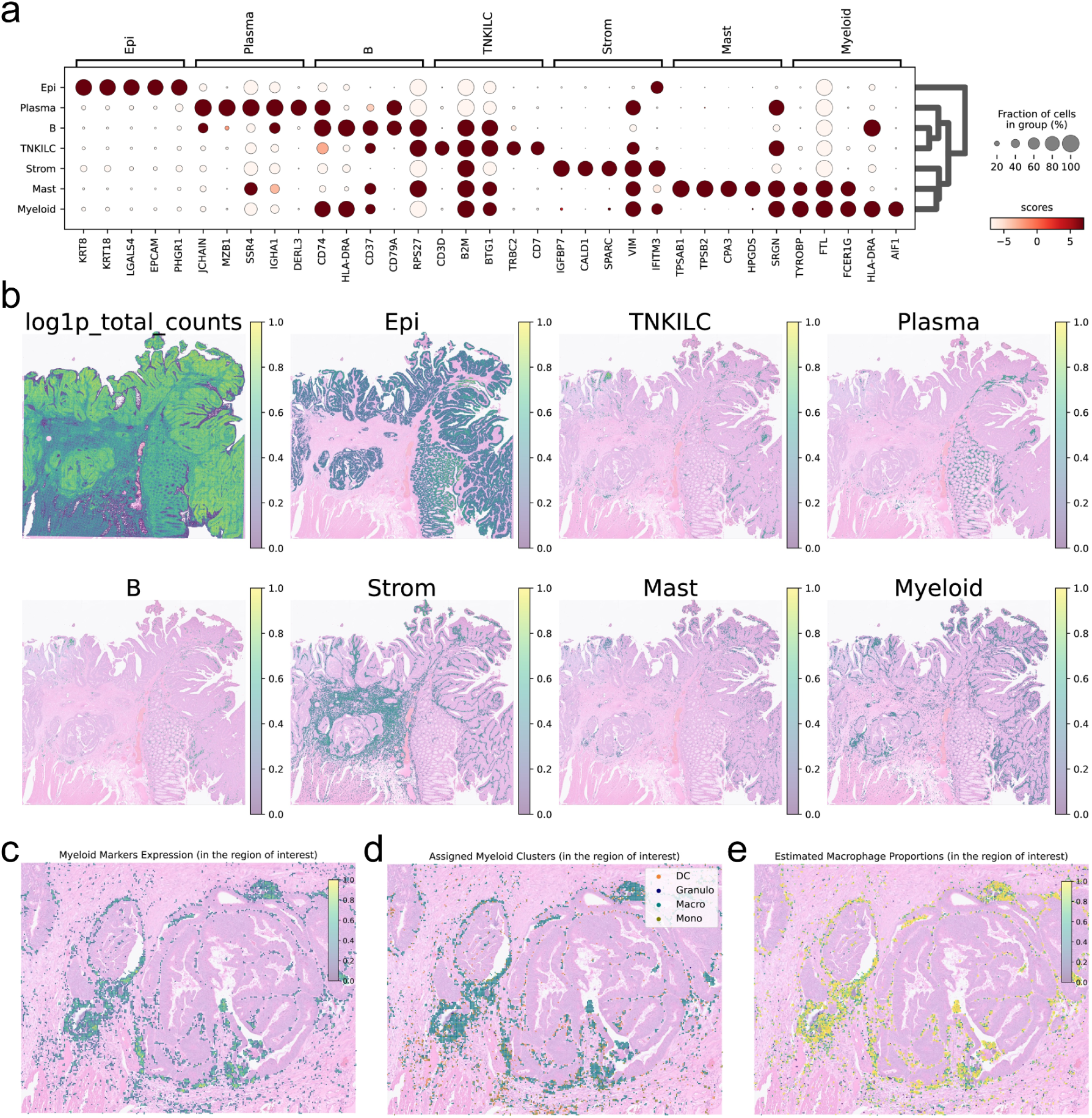
Easydecon uses global and local strategies to enable robust label transfer. **a,** The top 5 markers for each top-level cluster are shown as detected from the publicly available CRC dataset and we used 60 best markers overlapping with the ST object for the analyses. **b,** The top level cluster markers are visualized in each panel with the H&E tissue image at the background. The first panel shows the normalized total counts. We summed normalized expressions of the markers to aggregate them for each bin. Expression hotspots signal most likely locations of the clusters based on the permutation test (*p* < 0.01). **c,** ROI focused for myeloid cell markers shows the myeloid marker expression at this region. **d,** We assigned mid-level markers to myeloid hotspots showing that macrophages are the most common cell types. **e,** Cell type proportion estimates also confirm this.

In the second step, the left-over bins are used to refine granularity. An algorithm then measures cell type similarity to each bin. Because low-expression bins were already discarded, this stage runs with minimal noise and improved efficiency. Using the CRC dataset, we demonstrated this approach with myeloid cells, which comprise four main subtypes: dendritic cells (DC), granulocytes (granulo), macrophages (macro), and monocytes (mono). Thus, we increased the granularity by transferring these labels to myeloid cell bins (Fig. 2c,d) on a region of interest (ROI) containing a CRC tumor. In this example, the majority of bins are assigned to macrophages. It is also possible to infer cell type proportions. We predicted that the myeloid associated bins mostly contain macrophages (Fig. 2e).

### Easydecon can transfer cell types between multiple VisiumHD samples

Easydecon also allows the transfer of cluster information across multiple Visium HD samples. This workflow is similar to single-cell to spatial mapping. The source (or donor) spatial dataset (Sample P2) can be used to generate a DE table using standard single-cell pipelines such as Scanpy [17] or Seurat [18]. This table then can be transferred to another compatible spatial sample, enabling rapid analysis.

We tested this approach on two publicly available VisiumHD datasets from colon tissue, one CRC sample (Sample P5) and one normal tissue sample (Sample P3). Tissue clusters were taken from the official 10x Genomics object (Sample P2). After generating the DE table using these putative clusters, we transferred these to the other two samples. In this example, we applied only the similarity approach (i.e. phase two or high granularity step) and assigned each spot to the most similar cluster.

We included all clusters but visualized four key ones: two tumor-associated clusters plus the normal epithelial and smooth-muscle clusters (Fig. 3a). In the sample containing both tumor tissue and adjacent normal tissue, Easydecon clearly separated the tumor clusters from the normal epithelium and smooth muscle (Fig. 3b). In the purely normal colon sample, only the normal epithelial and smooth-muscle clusters were detected (Fig. 3c). These findings demonstrate that a single DE table from the donor slide is sufficient to map both CRC tumor-specific and normal tissue clusters to their correct spatial locations in other samples.

**Figure 3.**
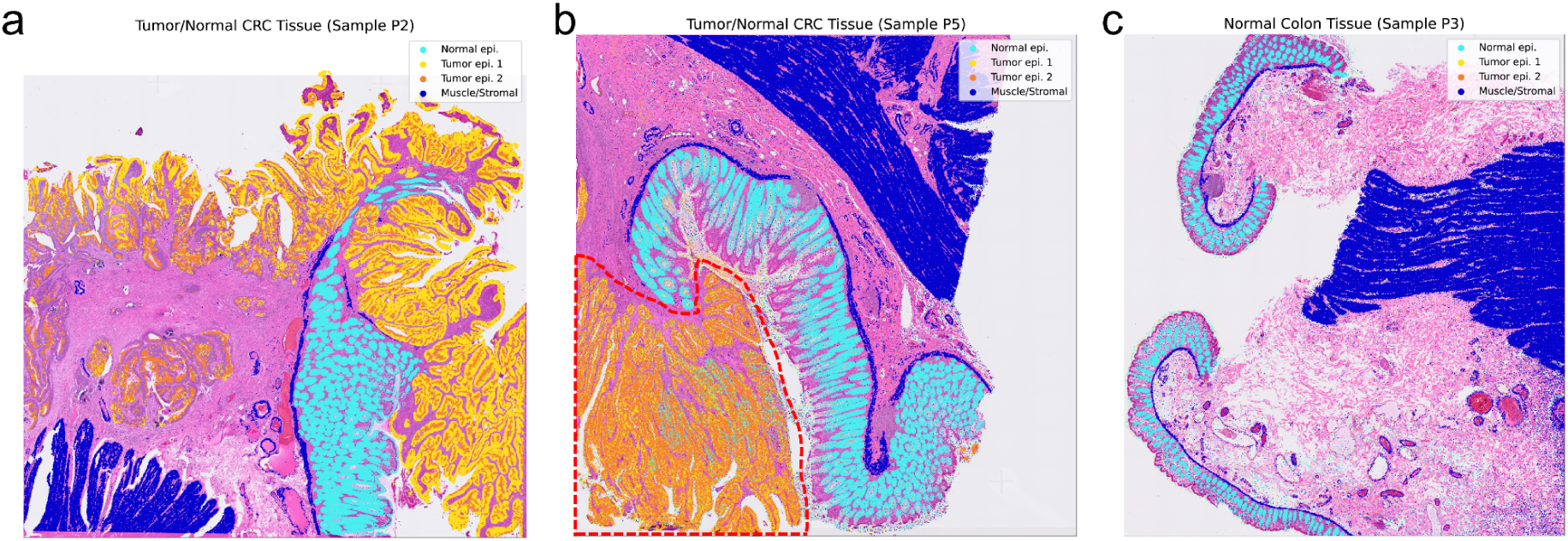
Spatial clusters are transferred to samples containing tumor and normal tissue. **a,** Clusters from tumor yellow/orange) and normal (cyan) epithelial regions were indicated. Smooth muscle cells were also indicated (blue). Note smooth muscle below normal epithelium (muscularis mucosae) and around vessels. **b,** The same regions are shown when cluster information is transferred from Sample 2 to Sample 5. Tumor cell area is indicated (broken line) **c,** Normal colonic tissue only contains the normal epithelial cluster and smooth muscle tissue. Even though we included tumor clusters, only normal tissue clusters were spatially transferred.

### Easydecon can be used with cell segmentation

The H&E tissue images of VisiumHD samples can be used for cell segmentation. Later, the segmented cells can be used with Easydecon for cell type transfer. Bin2cell is a computational method designed to reconstruct biologically meaningful single-cell transcriptomic profiles from high-resolution spatial transcriptomics data [13]. The tool aggregates transcriptomic signals from 2 µm bins into cellular units using morphological segmentation with the StarDist algorithm [19]. This bin-to-cell transformation enhances the accuracy of cell type annotation, provides higher per-cell gene detection and significantly improves spatial localization of cell populations compared to default analysis approaches using 8 µm bins. Bin2cell achieves these improvements efficiently, without the need for GPU computing resources which makes it an ideal tool to use with lightweight Easydecon.

To evaluate Bin2cell in conjunction with Easydecon, we selected another ROI in the Sample P5 spanning both tumor and normal tissue (Fig. 4a). Bin2cell aggregates 2 µm bins into segmented cells (Fig. 4b), which we then annotated using Easydecon and the same spatial clusters (Fig. 4c). Both the segmented and non-segmented analyses (on different resolutions) yielded consistent cell-type assignments, accurately separating normal and tumor regions (Fig. 4d,e,f). We also estimated cell-type proportions within the segmented cells using three different algorithms (Fig. 5a,b,c), and these results were highly concordant with the bin-based cell-type assignments.

**Figure 4.**
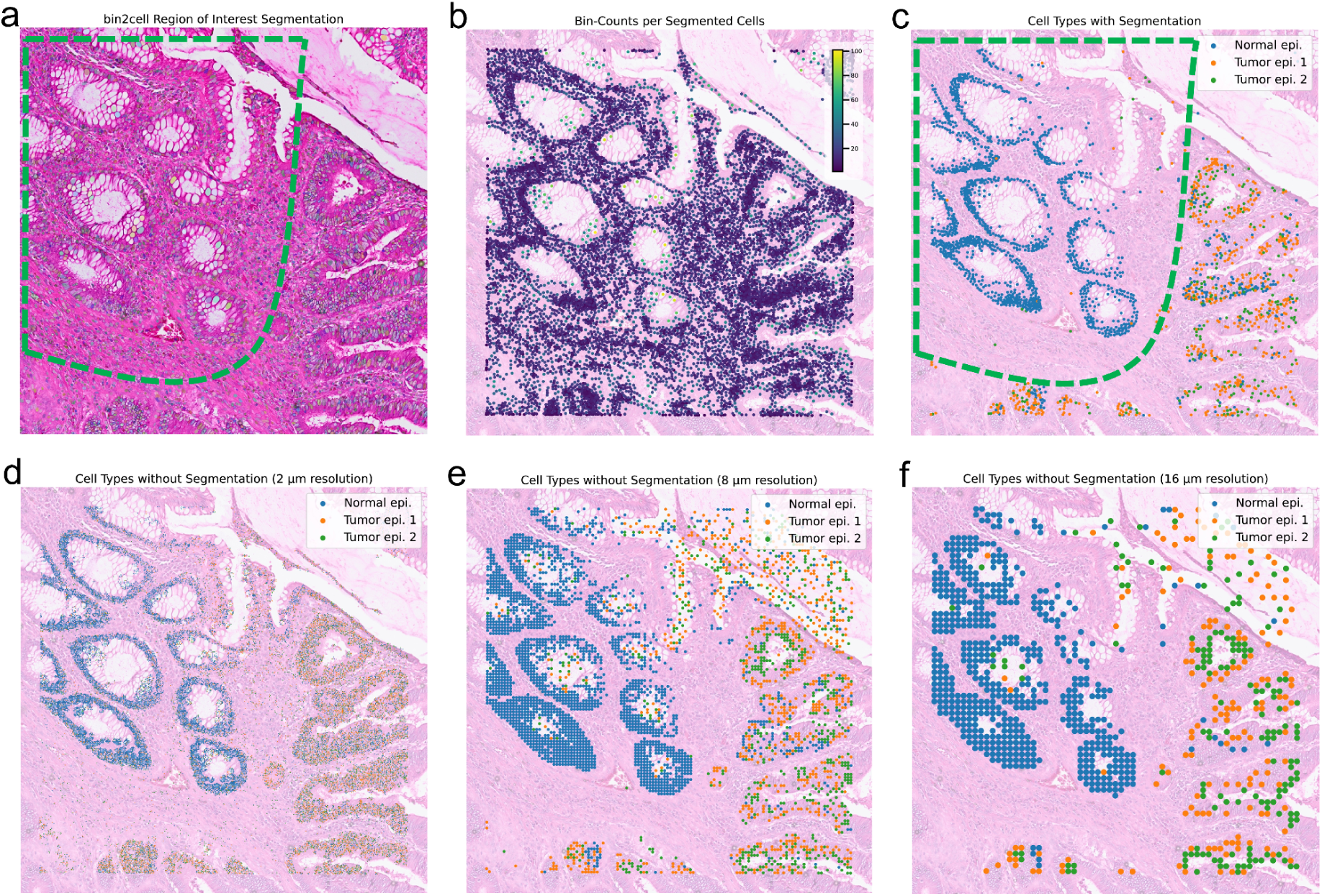
Easydecon can annotate segmented cells on human colon tissue. **a,** The H&E-stained bright-field image shows the selected ROI in Sample P5, encompassing both normal and tumor regions (the dashed outline indicates normal epithelial area). Segmentation within this ROI was performed using the Bin2cell tool. **b,** Segmented cells are color-coded according to the total number of 2 µm bins they contain, with higher values indicating more bins per that segment. **c,** Easydecon assigns cell types to the segmented cells via similarity-based label transfer, generating a precise spatial distribution of identified cell populations and clearly distinguishing normal from tumor tissue. **d, e, f** The ROI annotations without segmentation on different resolutions (2 µm, 8 µm and 16 µm) largely mirror the segmented results, underscoring Easydecon’s consistency and accuracy in both approaches.

**Figure 5.**
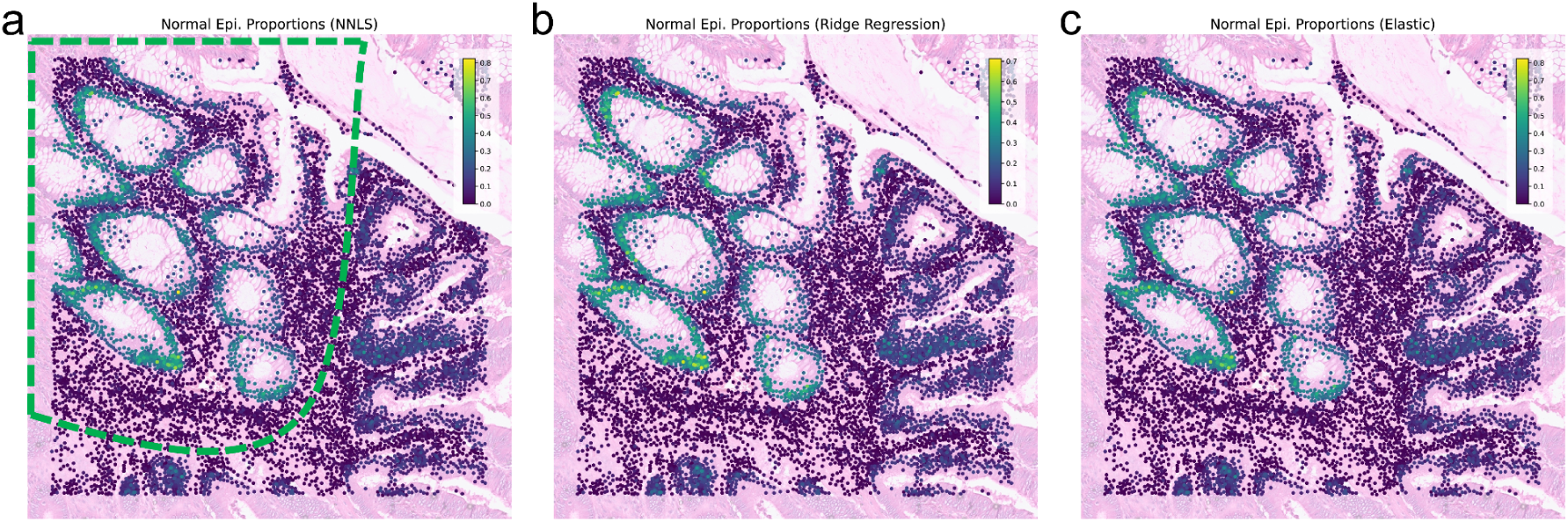
Cell type proportion estimates of segmented cells shows separation between normal and tumor tissues. **a,** Each spot represents a cell segmented by Bin2cell, with normal epithelial cluster proportions estimated via NNLS. The highlighted region corresponds to tumor-adjacent normal tissue. **b,** Ridge regression produces a similar distribution of proportions. **c,** Elastic-net regression likewise reflects the same pattern, providing sharper estimates for certain bins.

### Easydecon helps to differentiate between regional clusters

Using scRNA-seq we recently showed that macrophages within normal colonic tissue consist of several functionally distinct subsets [20], and those macrophages in the upper mucosa are transcriptionally different from those in the submucosal and muscular layers underneath. Building on this knowledge, we applied Easydecon to estimate tissue stratification within compiled single-cell macrophage datasets from multiple studies focused on tumor-adjacent regions [16,20–24], integrating them into a single object.

Adopting Easydecon’s two-phase approach, we initially identified macrophage expression hotspots (Fig. 6a) on normal colon tissue (Sample P3), which were predominantly located in the mucosal region. In the second phase, we transferred the annotated clusters to the hotspots identified (Fig. 6b,c) based on the macrophage clusters (Fig. 6d). Spatial analysis showed that LYVE1+ macrophages, in contrast to the other clusters, were primarily localized to the submucosa and muscularis layers (Fig. 6c). This was confirmed with unbiased neighborhood enrichment and co-occurrence analysis showing that LYVE1+ macrophages were spatially distinct and exhibited minimal interactions with other clusters (Fig. 6e,f).

**Figure 6.**
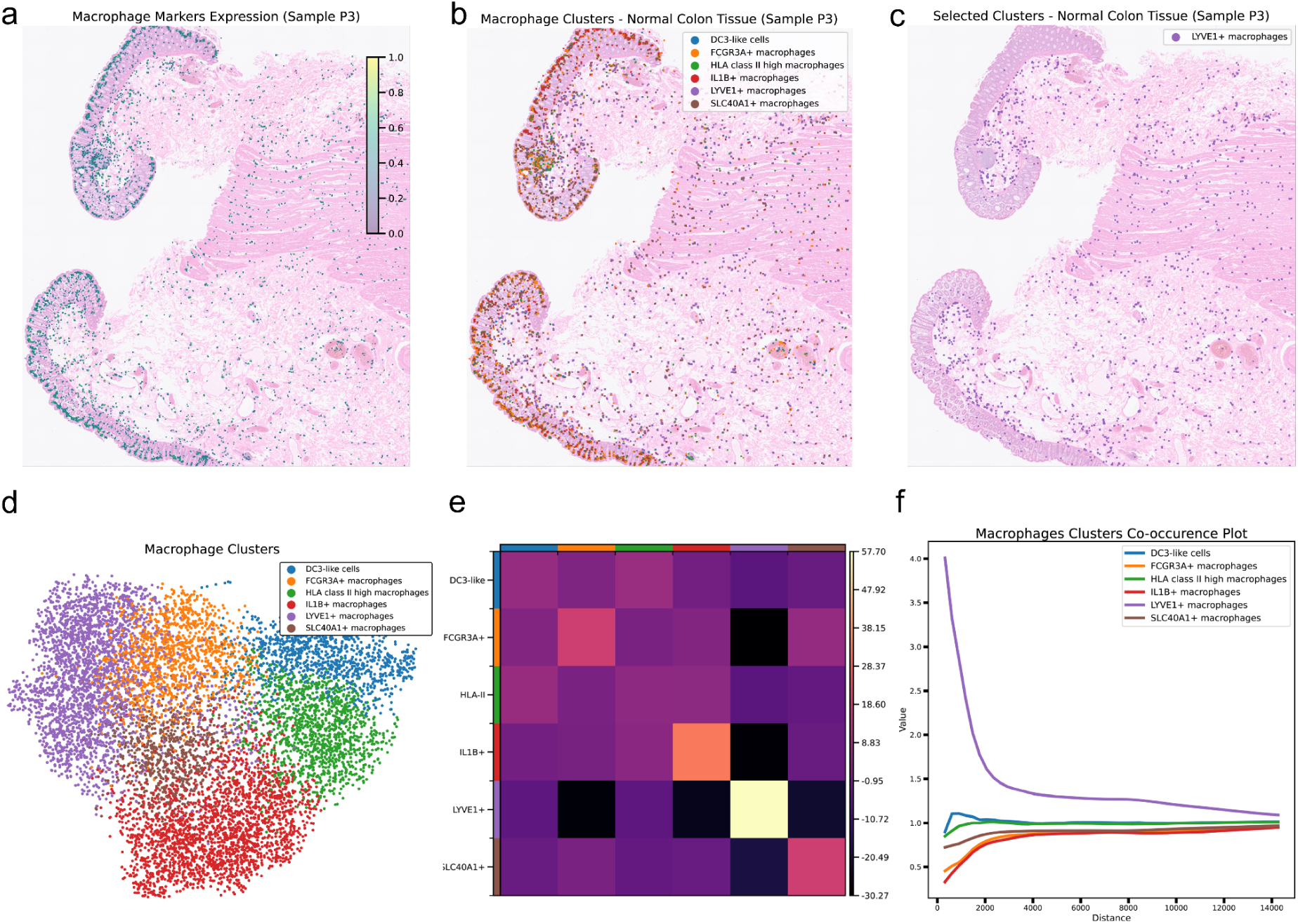
Easydecon can analyze spatial location specific macrophages. **a,** Macrophage expression hotspots, primarily located in the mucosal region. The scaling has been magnified for clarity. **b,** Easydecon transfers the macrophage clusters to the ROI. **c,** One specific cluster is highlighted: LYVE1+ macrophages, which are mainly found in the submucosa and muscularis region. **d,** The UMAP plot displays six macrophage clusters including DC-like cells. **e,** The neighborhood analysis shows if cells belonging to two different clusters/cell types are close to each other more often than expected. The results show that LYVE1+ cells are significantly distant from other mucosal cells, represented by darker colors. **f,** The co-occurrence plot also confirms this result. The purple line shows LYVE1+ macrophage cluster which is separated from other lines, representing other clusters.

### Easydecon performs better than other state-of-the-art methods

At the time of this study, most existing methods were not adapted for VisiumHD analysis, and many remain unsuitable. Thus, we compared Easydecon with four other compatible well-known methods, CellTypist [25], RCTD [26], CellAssign [27] and Tangram [28]. CellTypist focuses on automated cell type classification in scRNA-seq datasets, leveraging reference models and deep learning algorithms. While CellTypist excels at rapid identification of cell populations in single-cell level, its spatial functionality is limited. Yet, it is a recommended annotation method by Enact workflow [14]. CellTypist has a nice collection of reference models but it requires training to transfer other datasets. RCTD, on the other hand, provides a probabilistic framework specifically designed to deconvolve spatial transcriptomics data using single-cell references. Although RCTD accommodates multi-cell or partial overlap of transcripts per spot, it can be computationally demanding, especially at higher resolutions partially due to some of the restrictions of R programming. RCTD by default also utilizes 16 µm resolution to accommodate this. RCTD is the gold standard now and the best method for spatial deconvolution so far [10]. CellAssign is a probabilistic model designed to automatically assign single-cell data using predefined marker genes. It integrates biological prior knowledge with a statistical framework to compute probabilistic cell-type assignments [27]. Lastly, Tangram is a deep learning framework that maps single-cell RNA sequencing data onto spatial transcriptomics datasets coming from various platforms [28]. It is not officially supported for VisiumHD samples but we have included it as another method.

We tested all five methods using myeloid clusters on the ROI. All of them showed similar myeloid cell expression. However, Easydecon detected all myeloid subtypes available in colon cancer atlas (Fig. 7a). It is also by far the fastest method (Table 1) and most compatible. CellTypist missed all DCs both in the ROI and in the entire tissue (Fig. 7b). It also requires a RAW single-cell dataset. RCTD performed better than CellTypist. However, most of the cell types were still assigned to macrophages (Fig. 7c). Preprocessing of samples and more CPU time are also needed. CellAssign performed worse, missing most of the cell assignments. It also takes a vast amount of CPU time to run (Fig. 7d). Lastly, Tangram predicted more myeloid subtypes than the other methods (Fig. 7e) but its runtime is comparable to Easydecon if a GPU is used. To serve as a ground truth, we highlighted cell type markers showing macrophages (CD68, CD14), DCs (CD86, CD83, CD80), monocytes (S100A8-9, IL1B) and granulocytes (CD33, IL5RA, CCR3, MPO) (Fig. S1-S4). Easydecon detected cell types overlapping with ground truth marker expressions which are mostly missed by the others.

**Figure 7.**
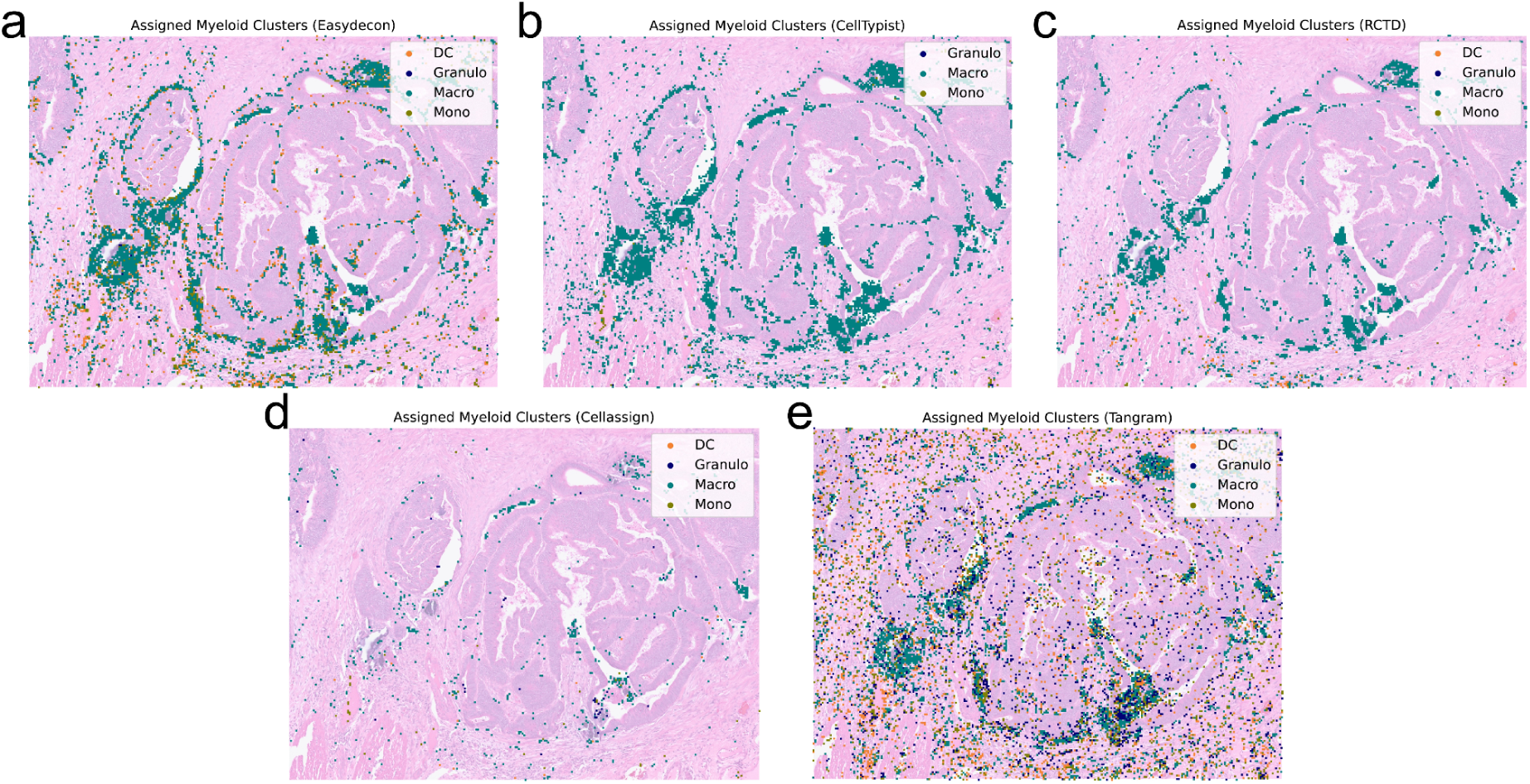
Myeloid cell type annotations obtained with five approaches. **a,** Easydecon recovers the complete myeloid repertoire present in the colon-cancer atlas, capturing DCs, macrophages, monocytes, and granulocytes. **b,** CellTypist labels most cells as macrophages and overlooks the DCs. **c,** RCTD similarly assigns the bulk of cells to macrophages, detecting only a handful of additional myeloid subtypes. **d,** CellAssign shows a similar pattern but it missed most of the myeloid subtypes. **e**, Tangram overpredicted myeloid subtypes compared to other methods but it still retains the same pattern.

**Table 1.** Summary of four spatial analysis methods.

|  | **Easydecon** | **CellTypist** | **RCTD** | **CellAssign** | **Tangram** |
| --- | --- | --- | --- | --- | --- |
| **Platform** | Python | Python | R | R | Python |
| **Scverse/Seurat Compatible** | Both (with DE table) | Only Scverse(Seurat to Anndata required) | Anndata to Seurat required | No (SCE object) | Only Scverse(Seurat to Anndata required) |
| **RAW data required** | No | Yes | Yes | No | Yes |
| **Training Required** | No | Yes | No | No | No |
| **GPU Required** | No | Yes* | No | Yes* | Yes* |
| **Runtime**<br>(Entire tissue 8 μm) | Preprocessing (DE table 1-5 mins)<br>Phase 1( 1-2 mins)<br>Phase 2 (5-30 mins) | Training (with CPU few hours)<br>Annotation (45 mins) | Preprocessing (10-30 mins)<br>Annotation (80 cores 3 hours) | 15 hours (80 cores, 500GB memory)** | Deconvolution (1-2 mins) |
| **System Used** | Laptop (M1) | HPC-GPU | HPC (80 cores) | HPC (80 cores) | HPC-GPU |

\* CPU can be used but heavily increases runtime. \*\* CPU was used for testing due to package compatibility issues.

For another test case, we downloaded publicly available VisiumHD tonsil dataset and human tonsil atlas single cell dataset. The data was processed as described in the material and methods section. We selected epithelial cells as a test case and used a set of compiled epithelial markers to detect hotspots. Later the epithelial clusters were transferred using the DE expression table (Fig. S5). The results showed clear separation between the most common epithelial cell types (i.e. basal and crypt cells) in the epithelial region [29], which is also successfully separated from the rest of the tonsil tissue.

## Discussion

There has been growing interest in scRNA-seq and ST due to their combined ability to reveal cellular identities and their spatial contexts within tissues [3,8]. Although various computational methods exist for integrating these modalities [12,14,17,18,30,31], the recent advancements introduced by VisiumHD require efficient and tailored analytical workflows. Here, we introduce Easydecon, a lightweight and user-friendly bioinformatics workflow specifically developed for rapid and reliable analysis of high-resolution VisiumHD spatial datasets using diverse scRNA-seq data sources.

A significant advantage of Easydecon is its foundational assumption that the 8 µm spatial bins provided by VisiumHD represent near-single-cell resolution and are thus homogeneous. This assumption effectively bypasses the need to handle complex scenarios, common in earlier versions of Visium technology. This simplifies downstream computational steps, enhancing analysis speed and reliability without compromising accuracy. In addition, Easydecon does not need exhaustive model training steps. Our head-to-head benchmarks confirm its speed and reliability compared with other popular methods. Easydecon surpassed them in terms of speed (Table 1) and cell type recovery. For example, Easydecon was able to detect overlooked myeloid cells (e.g. DC, monocytes) which are known to exist in the tumor and tumor microenvironment (TME) [32–34].

Easydecon addresses multiple critical aspects of spatial analysis, including marker visualization and cell type transfer. By employing robust similarity algorithms and permutation-based statistical approaches, Easydecon ensures precise identification of spatial expression hotspots. The flexible use of global and local markers allows researchers to integrate existing knowledge from literature or DE results from single-cell datasets seamlessly. However, the users do not need to employ the two phase approach but they can simply choose from hotspot detection or similarity approaches. We also included a wrapper so the entire workflow can be run in simple steps without much intervention, making it an easy and fast approach.

It is also possible to use Easydecon with available single-cell and spatial frameworks such as SpatialData [12], Squidpy [30] and Scanpy [17]. R language workflows like Seurat [18] are also supported via a DE table. Furthermore, Easydecon supports cell segmentation workflows, as demonstrated by our successful integration with Bin2cell [13]. This also does not break our foundational assumption (Fig. 4 and Fig. 5). Therefore, any available segmentation methods can be paired with Easydecon.

By aggregating high-resolution bins into biologically meaningful cellular units, Easydecon not only improves the spatial localization and accuracy of cell type annotations but also provides compatibility with downstream spatial analyses such as co-occurrence and neighborhood enrichment [30]. ST enables the identification of spatially relevant cell clusters, and Easydecon provides an efficient framework for their analysis. We tested Easydecon on a macrophage-focused single-cell dataset, demonstrating its capacity to distinguish macrophages derived from either the mucosal, submucosal or muscularis layers. Cluster distance analyses further confirmed this spatial partitioning (Fig. 6).

One limitation of Easydecon is its dependency on high-quality marker gene selection. Future work could explore automated and data-driven approaches for marker selection, potentially improving its accuracy and applicability across different biological contexts. It can be hard to decide a cut-off for a total number of markers to be used. This can also differ between phase one, expression hotspot detection, and phase two, granularity improvin. To overcome this, we recommend trying different parameters or methods that we include. For example, using different permutation parameters or testing different weight penalties in methods like weighted Jaccard could be a good starting strategy.

In addition, while Easydecon is optimized for the 8 µm bins commonly used in VisiumHD, extending its functionality to the higher-resolution 2 µm bins could further enhance its analytical power. If a smaller ROI is selected, it can benefit from increased accuracy while keeping the runtime relatively small. Selection of a ROI also helps with further parameter optimization. Easydecon can also benefit from a second filtering step depending on the cell types especially on crowded sections (Fig. 2c).

In summary, Easydecon offers a streamlined, efficient solution for analyzing complex spatial transcriptomics datasets, delivering rapid insights into both tissue biology and disease pathology. Its design circumvents the need for model training and specialized hardware, such as GPUs or high-performance computing environments, making it accessible to a wider range of laboratories and researchers with limited computational resources. By leveraging a two-phase approach and robust similarity algorithms, Easydecon seamlessly integrates single-cell and spatial data, enabling more precise and user-friendly analyses.

## Data availability

The CRC atlas dataset was downloaded using the accession SCP1162 [16]. We also used 3 Visium HD slides, two representing CRC patients and one representing normal colon. All are taken from http://10xgenomics.com/datasets. To perform region’s differentiation we used single cell data available from [16,20–24]. We shared the DE tables and sample files at our GitHub. The tonsil dataset is downloaded from BioConductor [15] and the tonsil VisiumHD dataset is from 10x genomics http://10xgenomics.com/datasets.

## Code availability

The source code of Easydecon is made available via GitHub, https://github.com/sinanugur/easydecon. All scripts used to reproduce the analyses included in this manuscript can be downloaded from the GitHub repository. Easydecon is also available at PyPi. All the development versions are available as a requirement file which is a part of Zenodo deposited file available at https://doi.org/10.5281/zenodo.17711754.

The supplementary file contains the supplementary figures.

## Acknowledgements

The work was made possible through generous funding by Norwegian Cancer Society (no. 223245 and 273170).

## Author contributions

SUU devised the project, created the workflow and Python scripts, software, and drafted the manuscript with input from all authors. VTK contributed to the data analyses, testing, contributed to the initial discussions and manuscript draft. ESB revised the manuscript and acquired financial support. FLJ revised the manuscript and acquired financial support. DD supervised the project, contributed to the workflow discussion, revised the manuscript and acquired financial support.

