## supplementary file for "Easydecon: Efficient Cell Type Mapping for High-Definition Spatial Transcriptomic Data"

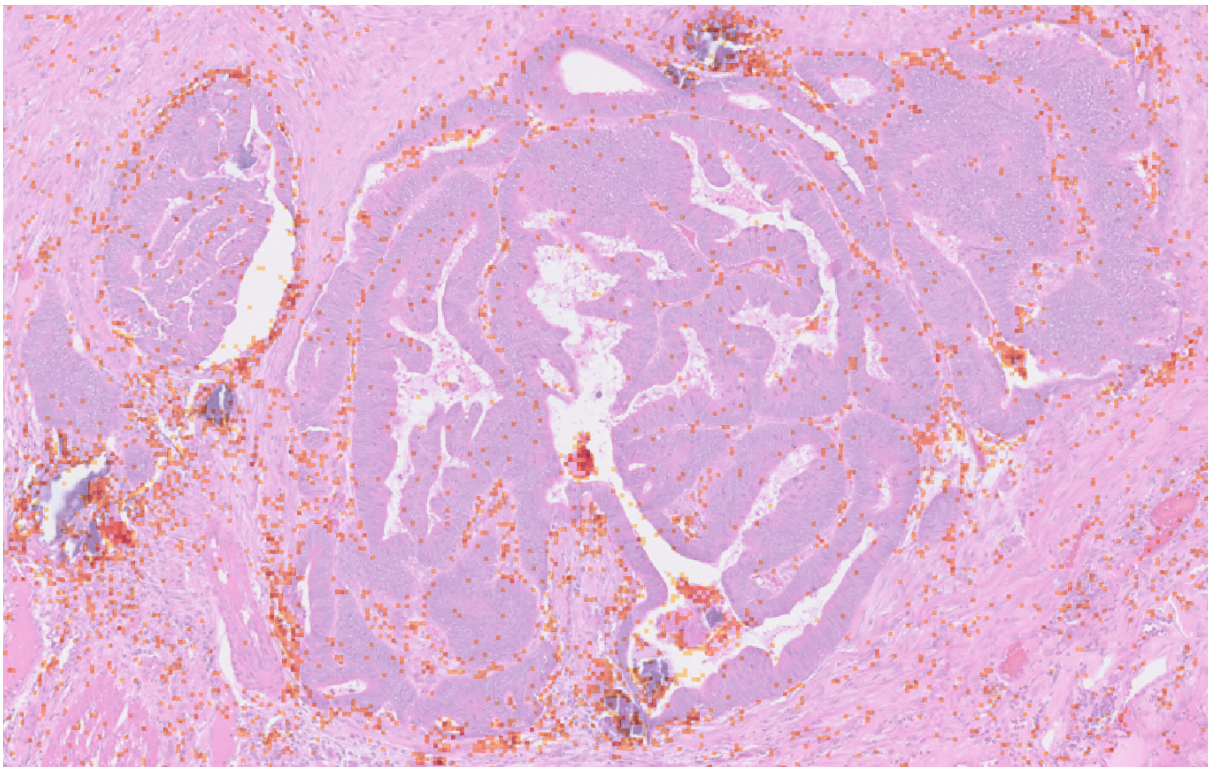

**Supplementary Figure 1.** CD68, CD14 macrophage markers summed log2 expression.

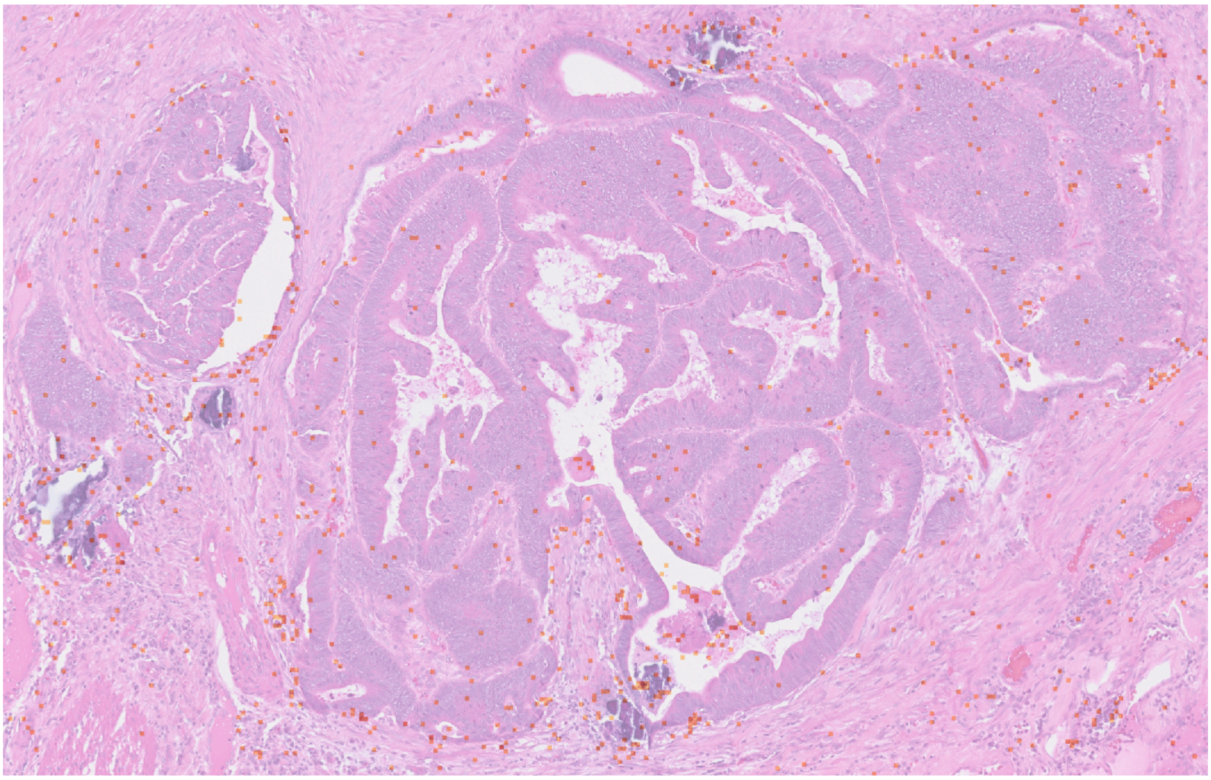

**Supplementary Figure 2.** CD86, CD83, CD80 mature dendritic cells (DC) markers summed log2 expression.

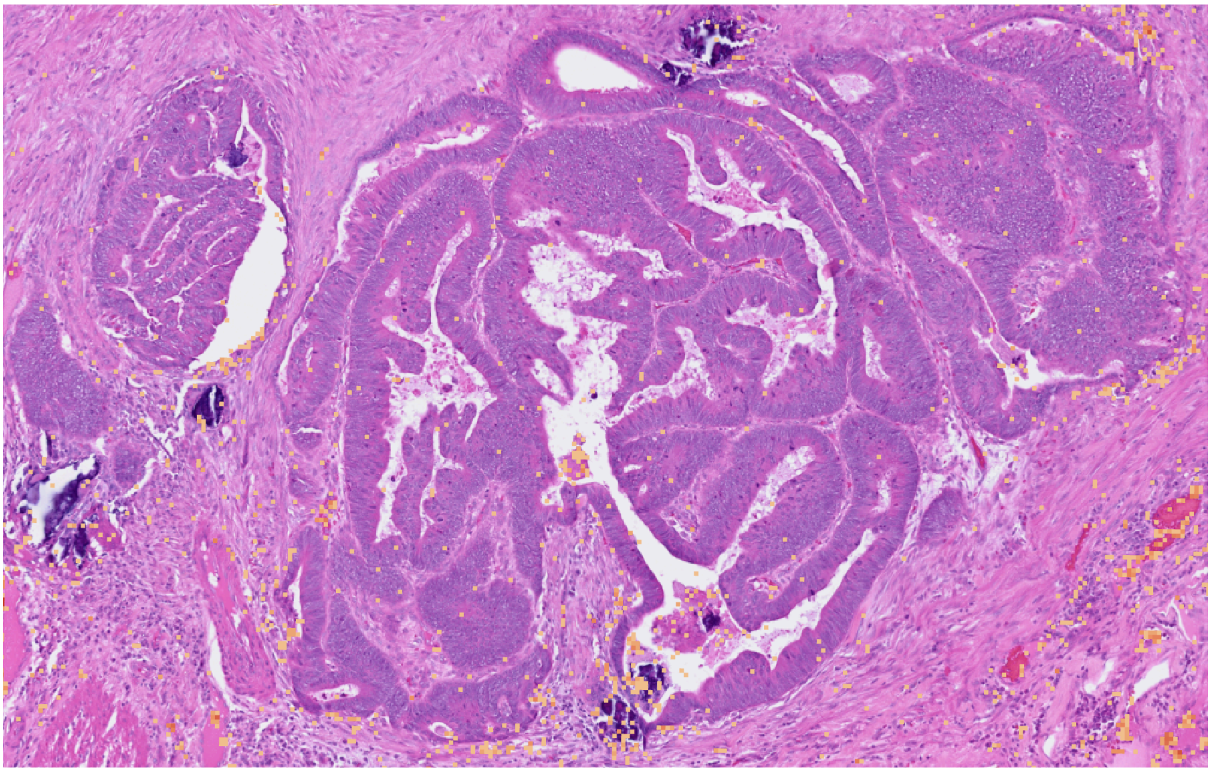

**Supplementary Figure 3.** S100A8-9 and IL1B monocyte markers summed log2 expression.

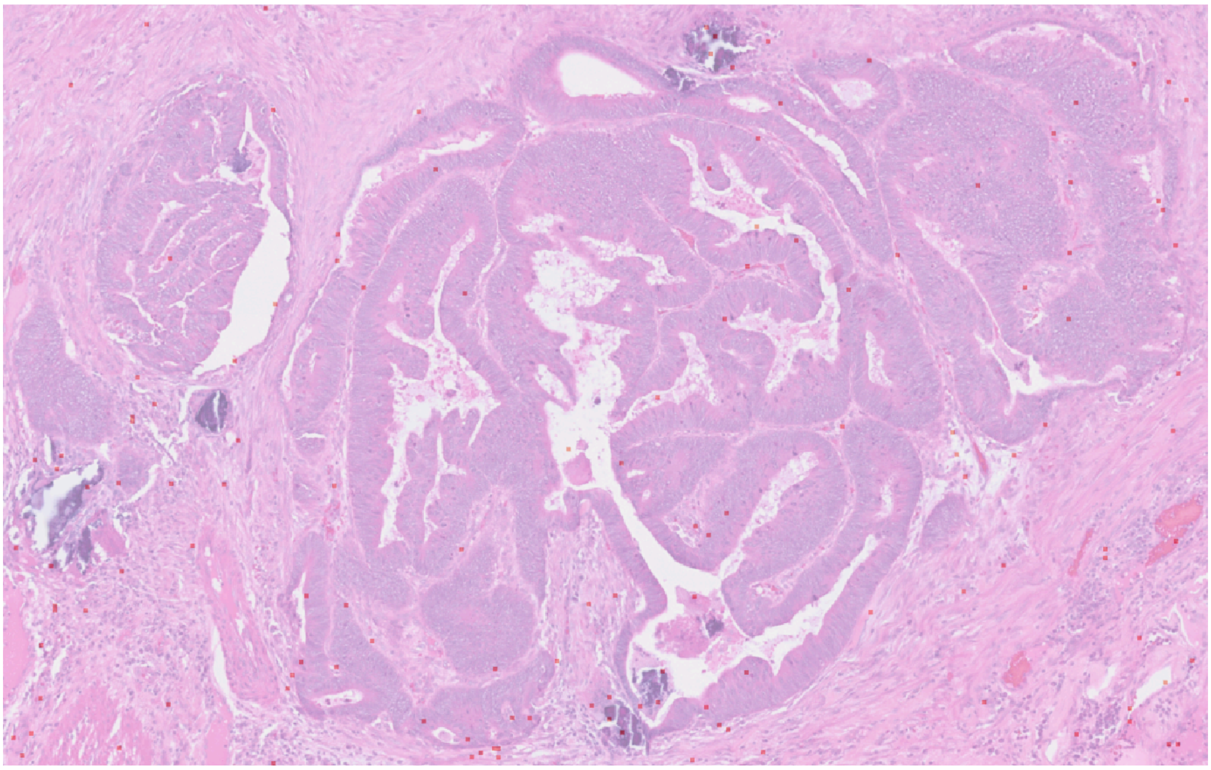

**Supplementary Figure 4.** CD33, IL5RA, CCR3 and MPO granulocytes markers summed log2 expression.

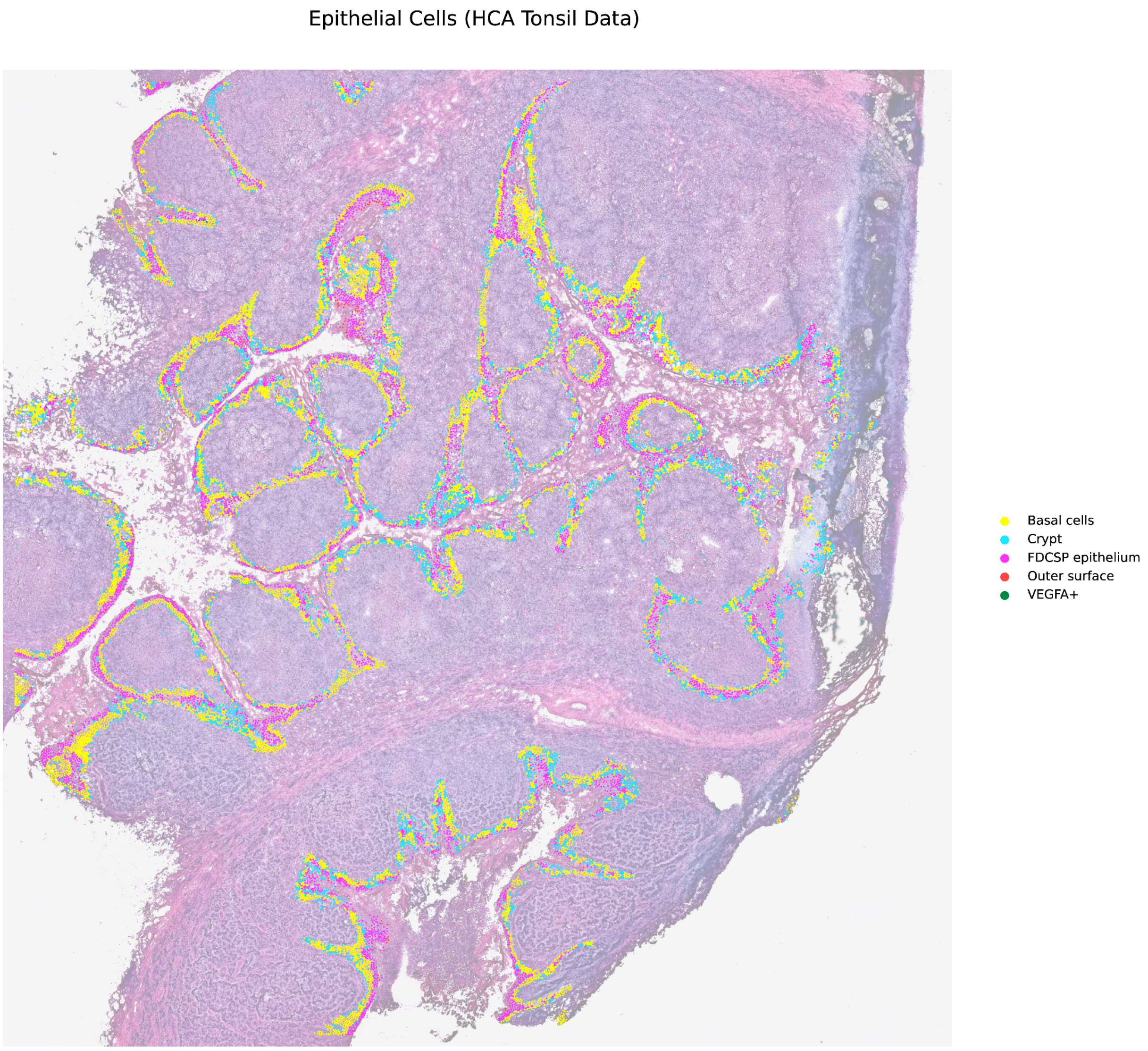


**Supplementary Figure 5.** We predicted the positions of tonsil epithelial cells in a VisiumHD dataset. The results showed clear epithelial patterns.
